# Profiling the intratumoral immune landscapes of primary and syngeneic Kras-driven sarcoma mouse models

**DOI:** 10.1101/2020.07.01.181263

**Authors:** Wade R. Gutierrez, Amanda Scherer, Gavin R. McGivney, Vickie Knepper-Adrian, Emily A. Laverty, Grace Roughton, Rebecca D. Dodd

**Author notes:** **Corresponding author:** Rebecca D. Dodd, Carver College of Medicine, University of Iowa, 285 Newton Rd, 3269C CBRB, Iowa City, Iowa 52246.

## Abstract

The introduction of immunotherapy has revolutionized cancer treatment and fueled interest in the immune cell composition of preclinical tumor models. Both genetically-engineered mouse models and syngeneic cell transplant approaches use immunocompetent mice to study mechanisms driving immunotherapy response and resistance. Due to their rapid and reproducible nature, there has been an expanded interest in developing new syngeneic tools from established primary tumor models. However, there are few analyses directly comparing the immune profiles of primary models with their derived syngeneic counterparts. Here we report comprehensive immunophenotyping of primary tumors from two well-established sarcoma models and four syngeneic allografts derived from these models. We observed that cell lines derived from the same type of genetically engineered primary tumor form allografts with significantly different levels of immune infiltration and intratumoral immune cell composition. Most notable are the differences in the T cell compartment of the syngeneic models, including CD4+ T cell, CD8+ T cell, and regulatory T cell populations – each of which have well-documented roles in tumor response to immunotherapy. Our findings highlight the importance of thorough characterization of syngeneic models prior to their use in preclinical studies in order to maintain scientific rigor and to increase the translatability of findings from bench to bedside.

## 1. INTRODUCTION

Mouse models are an important cornerstone of research for cancer therapeutics.^1,2^ These tools are used in preclinical proof-of-concept studies for novel therapeutic paradigms and provide an invaluable platform for mechanistic studies. Implantation of human tumor cell lines into immunocompromised mice (xenografts) are vital to understanding responses to traditional cytotoxic or targeted therapies. However, these models do not allow for study of the immune system and cannot be used to examine immunotherapy. In particular, the ability to model a tumor in an animal with an intact immune system facilitates study of tumor sensitivity, immune infiltration, and the importance of distinct immune cell subtypes in treatment response. Two approaches that utilize immunocompetent hosts are genetically-engineered mouse models (GEMMs) and syngeneic tumor models.^3,4^ In GEMMs, primary tumors are generated *de novo* from genetically-modified mice that contain mutations commonly found in human cancer. These tumors develop in a native, immune-proficient microenvironment. While these tumors can reflect the heterogeneity of human cancers, they can require large numbers of mice and extended latency periods. In contrast, syngeneic models (allografts) are generated by implanting murine cancer cells into mice of the same background strain as the original tumor. Syngeneic approaches allow for rapid and reproducible preclinical screens in animals with an intact immune system due to the short latency periods, relatively uniform tumor growth, and lower tumor heterogeneity. However, it is not known how well these syngeneic models reflect the immune composition of the primary GEMM models.

Recently, there have been several efforts to characterize the diverse immune cell profiles and immunotherapy responses across the plethora of syngeneic approaches. These studies have found a wide heterogeneity in total tumor immune infiltrates across syngeneic models^5–9^, ranging from immune-rich (~40% live cells in RENCA tumors) to immune-poor (~4% live cells in B16F10 tumors). Each syngeneic model also has a distinct composition of immune cells. For example, among BALB/c syngeneic models, MDSCs are the most abundant immune cells in 4T1 breast tumors, while macrophages are dominant in RENCA kidney tumors, and NK cells comprise the majority of immune cells in CT26 colorectal cancers.^5^ Even among the same syngeneic model, tumor location can alter the immune profile, as orthotopically injected CT26 tumors contain higher levels of T, B, and NK cells than tumors grown subcutaneously.^10^ However, there are few studies that examine how well syngeneic models reflect the immune profile of their original primary tumors. Ultimately, a better understanding of immune infiltrates between primary tumors and their syngeneic counterparts will help prioritize selection of these models and aid in extrapolating their data to the clinic.

Here, we describe a comprehensive immunological characterization of a new syngeneic cell series and their parental primary tumors from murine soft-tissue sarcomas, aggressive tumors of the connective tissue. Our group generated several syngeneic tumor models from two well-characterized primary mouse models soft tissue sarcoma driven by Kras activation and p53 loss.^11–16^ The first develops high-grade undifferentiated pleomorphic sarcoma (UPS) that resembles human tumors at the molecular, histologic, and physiologic levels. Injection of adenovirus expressing Cre recombinase (Ad-Cre) into the gastrocnemius muscle of LSL-Kras^G12D^; Trp53^Flox/Flox^ mice (KP mice) results in activation of oncogenic Kras^G12D^ and deletion of Trp53. Approximately 10 weeks after Ad-Cre injection, tumors develop at the site of injection. The second develops rhabdomyosarcoma (RMS), the most common pediatric soft-tissue sarcoma. Injection of 4-hydroxytamoxifen (TMX) into the gastrocnemius muscle of Pax7-Cre-ER; LSL-Kras^G12D^; Trp53^Flox/Flox^ mice (Pax7KP mice) results in activation of Kras^G12D^ and deletion of Trp53 in muscle satellite cells. Tumors develop at the site of injection approximately 6 weeks after TMX injection. We established multiple cell lines from these tumors that we have termed them <u>K-R</u>as <u>I</u>nduced <u>M</u>urine <u>S</u>arcoma lines (KRIMS). We orthotopically implanted KP KRIMS cells into 129/SvJae mice to generate syngeneic UPS models and P7KP KRIMS cells into P7KP mice to generate syngeneic RMS models. Using flow cytometry, we characterized the immune cell infiltrate of both the primary and syngeneic UPS and RMS tumors. This study provides important characterization of the intratumoral and systemic immune profiles between primary and syngeneic tumor models, which has strong implications for future preclinical immunotherapy studies using murine approaches.

## 2. METHODS

### Mice

The LSL-Kras^G12D^, Trp53^Flox/Flox^ (KP) mouse tumor model was previously described.^11^ Briefly, Ad-Cre (University of Iowa Viral Vector Core, Iowa City, Iowa) was mixed with calcium phosphate, incubated for 15 minutes at room temperature, and injected into the gastrocnemius muscle (50 μL). For the P7KP RMS model, Pax7-Cre-ER; LSL-Kras^G12D^; Trp53^Flox/Flox^ mice were injected with 4-hydroxytamoxifen (Sigma-Aldrich, H7904) in the gastrocnemius muscle as previously described.^17^ Tumors were measured by digital caliper three times weekly once tumors reached 150-275 mm^3^, and volume was calculated using the formula V = (π x *L* x *W* x *H*)/6, with *L*, *W*, and *H* representing the length, width, and height of the tumor in mm, respectively. Male and female mice were used for all studies. All animal experiments were performed in accordance with protocols approved by IACUC at the University of Iowa.

### Derivation of cell lines

Cell lines were derived from terminally-harvested tumors. A portion of tumor tissue was stored in 10% neutral buffered formalin for fixation and subsequent paraffin embedment. Remaining tumor tissue was dissected using forceps and surgical scissors, washed in 5 mL PBS in a 6-well plate, then finely minced using surgical scissors. 5 mL of Dissociation Buffer (Collagenase Type IV (700 units/mL, Gibco, 17104-019) and dispase (65 mg/mL, Gibco, 17105-041) in PBS) was added to each well. Plates were incubated for one hour at 37°C on an orbital shaker and transferred to a tissue culture hood. Dissociated tissue was passed through a sterile 70 μM cell strainer (Fisherbrand, 22363548) into a 50 mL conical vial using a 10 mL serological pipette and the plunger from a 1 mL syringe (BD, 309628). Cell strainers were washed with 25 mL sterile PBS into corresponding conical vials. Cell suspensions were centrifuged and cell pellets were resuspended and plated in DMEM (Gibco, 11965-092). Cells were grown in 10 cm dishes maintained in DMEM media containing 10% FBS, 1% penicillinstreptomycin (Gibco, 15140-122) and 1% sodium pyruvate (Gibco, 11360-070). When 90% confluency was reached, 15-35% of cells were passaged into a new dish. After a minimum of 10 passages, cells were frozen down to be used for syngeneic injections.

### Syngeneic allografts

All cells were ~ 90% confluent on the day of injection. Cells were trypsinized with 0.25% trypsin, washed, and resuspended in sterile PBS containing calcium chloride and magnesium chloride. Three-to six-month-old 129/SvJae or P7KP mice were injected with 50 μL of cell suspension in the left gastrocnemius muscle using a 31G needle. For UPS allografts, 129/SvJae mice were injected with either KRIMS-1 (2.5×10^5^ cells) or KRIMS-2 (1×10^6^ cells). For RMS allografts, P7KP mice were injected with either KRIMS-3 (2×10^5^ cells) or KRIMS-4 (5×10^4^ or 1×10^5^ cells). Once an initiation volume of 150-275 mm^3^ was reached, tumors were measured three times weekly by digital caliper.

### Histological analysis

Formalin-fixed paraffin embedded tumors were sectioned and stained with hematoxylin (Vector Laboratories, H-3401) and eosin (Harleco, 200-12) to evaluate tissue morphology. Images were taken using an Olympus BX61 microscope (Olympus) at 40x magnification.

### Flow cytometry immunoprofiling

Tissue from terminally-harvested tumors was washed in 5 mL PBS in a 6-well plate, then finely minced using surgical scissors. 4.5 mL Collagenase Type IV (700 units/mL, Gibco, 17104-019) and 0.5 mL FBS were added to each well containing tumor tissue. Plates were incubated for one hour at 37°C on an orbital shaker. Following incubation, dissociated tissue was passed through a 70 μM cell strainer into a 50 mL conical vial using a 10 mL serological pipette and the plunger from a 1 mL syringe. Cell strainers were washed with 25 mL PBS into corresponding conical vials. Spleens from mice were finely minced with scissors, then passed through a 70 μM cell strainer without enzymatic dissociation. Cell suspensions were centrifuged and cell pellets were resuspended in 2 mL ACK lysis buffer (Gibco, A1049201). After five minutes, 10 mL PBS were added and samples transferred to 15 mL conical tubes and centrifuged at 500xg for 5 minutes. Cell pellets were resuspended in cell staining buffer (Biolegend, 420201). In a round-bottom 96-well plate, 50 μL aliquots of cell suspensions were incubated with anti-CD16/32 (clone 93, Biolegend) to block Fc receptors and Zombie Aqua Viability Dye (Biolegend, 77143) on ice. After a 10-minute incubation, 50 μL of antibodies were added and incubated on ice for 30 min. Antibodies used were as follows: anti-CD45 BV605 (clone 30-F11, Biolegend), anti-CD11b PE (clone M1-70, Biolegend), anti-CD11c

BV421 (clone N418, Biolegend), anti-CD3 PE-Cy7 (clone 145-2C11, Biolegend), anti-CD4 Alexa Fluor 700 (clone GK1.5, eBioscience), anti-CD8 PerCP/Cy5.5 (clone 53-6.7, Biolegend), anti-CD25 PE-Cy5 (clone PC61.5, Invitrogen), anti-NKp46 PerCP/Cy5.5 (clone 29A1.4, Biolegend), anti-F4/80 Alexa Fluor 488 (clone BM8, Biolegend), anti-B220 APC (clone RA3-6B2, Biolegend), and anti-I-A/I-E Alexa Fluor 700 (clone M5/114.15.2, Biolegend). UPS primary tumors and spleens and naïve KP spleens were stained for regulatory T cells using anti-FoxP3 APC (clone FJK-16s, Invitrogen) and the FoxP3/transcription factor staining buffer set (eBioscience, 00-5523-00) according to the manufacturer’s instructions. P7KP primary tumors and spleens; KRIMS allograft tumors and spleens; and naïve 129/SvJae spleens were stained for regulatory T cells with anti-CD127 APC (clone SB/199, Biolegend). Cells were fixed with fix buffer (Biolegend, 420801) and stored in the dark at 4°C for 24-48 hours. Samples were analyzed with a BD LSR II flow cytometer. Data analysis was performed using FlowJo version 10.6.1 (Becton, Dickinson and Company). Fluorescence minus one (FMO) controls were used to set the boundary gates between positive and negative populations.

### Statistical analysis

Statistical analysis was performed using GraphPad Prism 8. Tumor time to triple was analyzed using Welch’s ANOVA. Immune profiles were analyzed using either unpaired t tests with Welch’s correction or Welch’s ANOVA.

## 3. RESULTS

### 3.1 Comparison of genetically engineered UPS and RMS primary tumors

UPS and RMS primary tumors were induced in the gastrocnemius muscle KP and Pax7KP mice, respectively, as previously described.^11–17^ Both models form soft tissue sarcomas and rely on expression of oncogenic *Kras^G12D^* and deletion of *Trp53* to drive tumor formation. The Pax7 promoter for Cre-ER limits *Kras^G12D^* expression and *Trp53* deletion to muscle satellite cells, the RMS cell of origin. For UPS, the exact cell of origin remains unknown. Though the models rely on the same initiating mutations and are induced at the same an atomic location, the time to tumor formation differed greatly. UPS primary tumors developed 9-17 weeks after AdCre injection (average 12.4 weeks). In contrast, RMS primary tumors developed significantly faster, initiating 5-7 weeks after TMX injection (average 6.0 weeks) (Figure 1A). Differences in tumor initiation did not correlate with differences in tumor growth rates or CD45+ immune cell infiltration (Figure 1B and 1C). In fact, proportions of several immune cell subsets including T cells, B cells, tumor associated macrophages (TAMs), NK cells, dendritic cells, and monocytes/polymorphonuclear cells (PMNs) were almost identical in UPS and RMS primary tumors (Figure 1D). However, within the T cell compartment, we found significant differences in levels of CD4+ T cells and CD8+ T cells (Figure 1E). UPS primary tumors contained a significantly higher proportion of CD8+ T cells and a significantly lower proportion of CD4+ T cells than the RMS primary tumors. No differences in regulatory T cells (Tregs) were observed. These findings highlight T cell-specific alterations to the intratumoral immune landscape driven by tumor cell of origin rather than genetic alterations or anatomic location.

**Figure 1:**
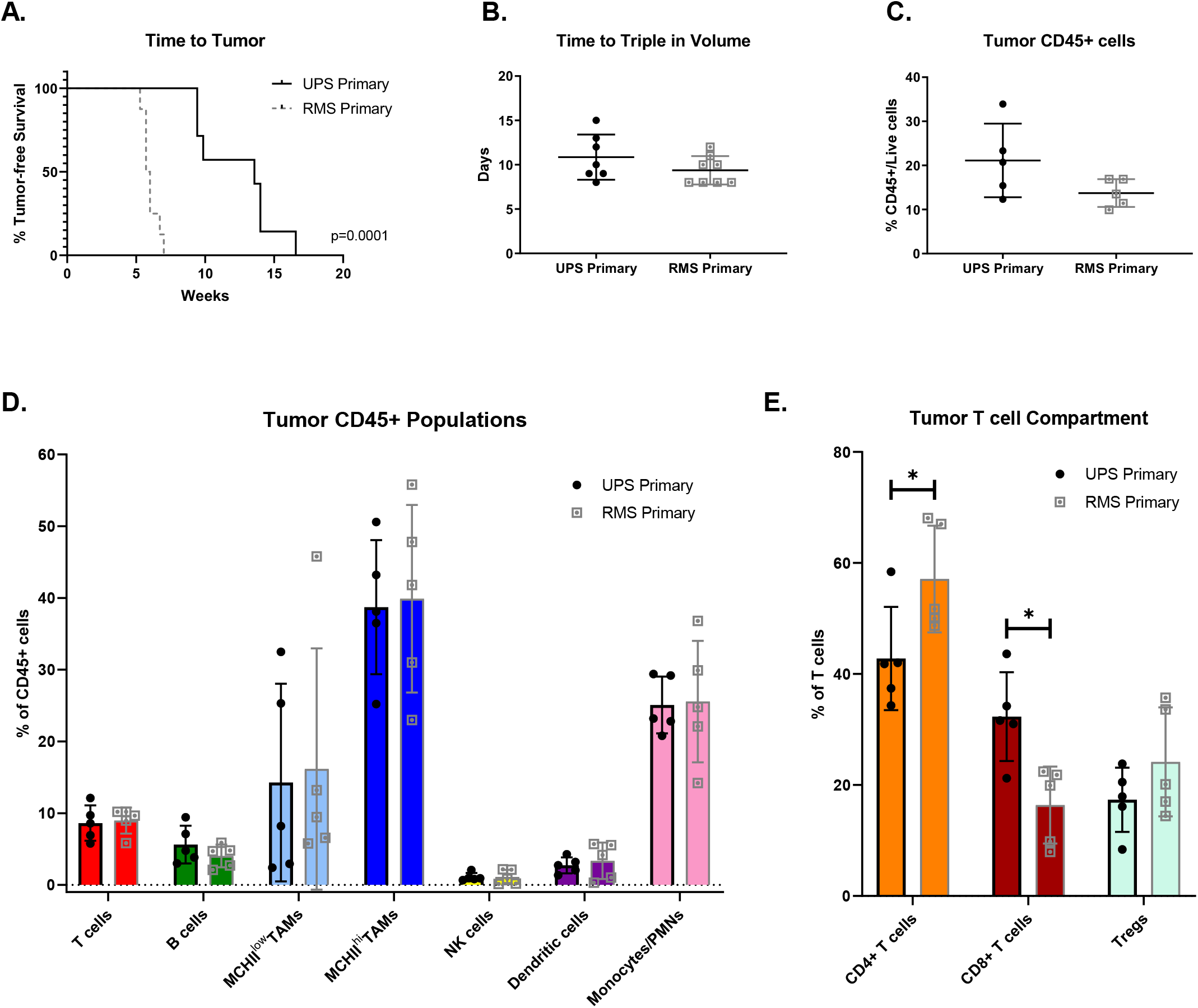
UPS and RMS primary tumors differ in initiation time and intratumoral T cell compartments. (A) Time to tumor initiation following injection of Ad-Cre (UPS) or TMX (RMS). (B) UPS and RMS primary tumors grow at similar rates, tripling in volume after approximately 10 days. (C) CD45+ cell infiltration is similar in UPS and RMS primary tumors. (D) Levels of several immune cell subtypes do not differ between the two models. (E) Significant differences between UPS and RMS primary tumors are evident within the T cell compartment.

### 3.2 Generation of syngeneic sarcoma models from primary tumors

Next, we generated four orthotopic allograft models of UPS and RMS using KRIMS cell lines derived from primary tumors. To generate cell lines, we enzymatically digested individual primary tumors from KP and Pax7KP mice (Figure 2A). Each tumor was passaged 10 times to remove stroma and establish a cell line. Based on *in vitro* growth phenotypes, we chose KRIMS-1 and KRIMS-2 (UPS) and KRIMS-3 and KRIMS-4 (RMS) for further *in vivo* characterization. Syngeneic tumors developed 1.5 to 3 weeks after injection of cells into the gastrocnemius muscle and showed growth rates similar to their primary tumor counterparts (Figure 2B and 2D). KRIMS-3 syngeneic tumors showed a slight, but statistically significant, reduction in tumor tripling time (6.3 days) compared to RMS primary tumors (9.4 days), while KRIMS-4 syngeneic tumors grew more similarly to RMS primary tumors with a tumor tripling time of 8.0 days (Figure 2D). Histological analysis confirmed that KRIMS tumors had pathological features similar to the primary tumors. UPS primary and syngeneic tumors contained multiple mitoses, enlarged nuclei, and spindle cell morphology (Figure 2C). RMS primary and syngeneic tumors contain small, round cells with hypochromic nuclei and a more eosinophilic cytoplasm (Figure 2E). Taken together, the similar growth rates and pathological features demonstrate that the syngeneic models have a high-fidelity to the primary model and support their utility as preclinical platforms.

**Figure 2:**
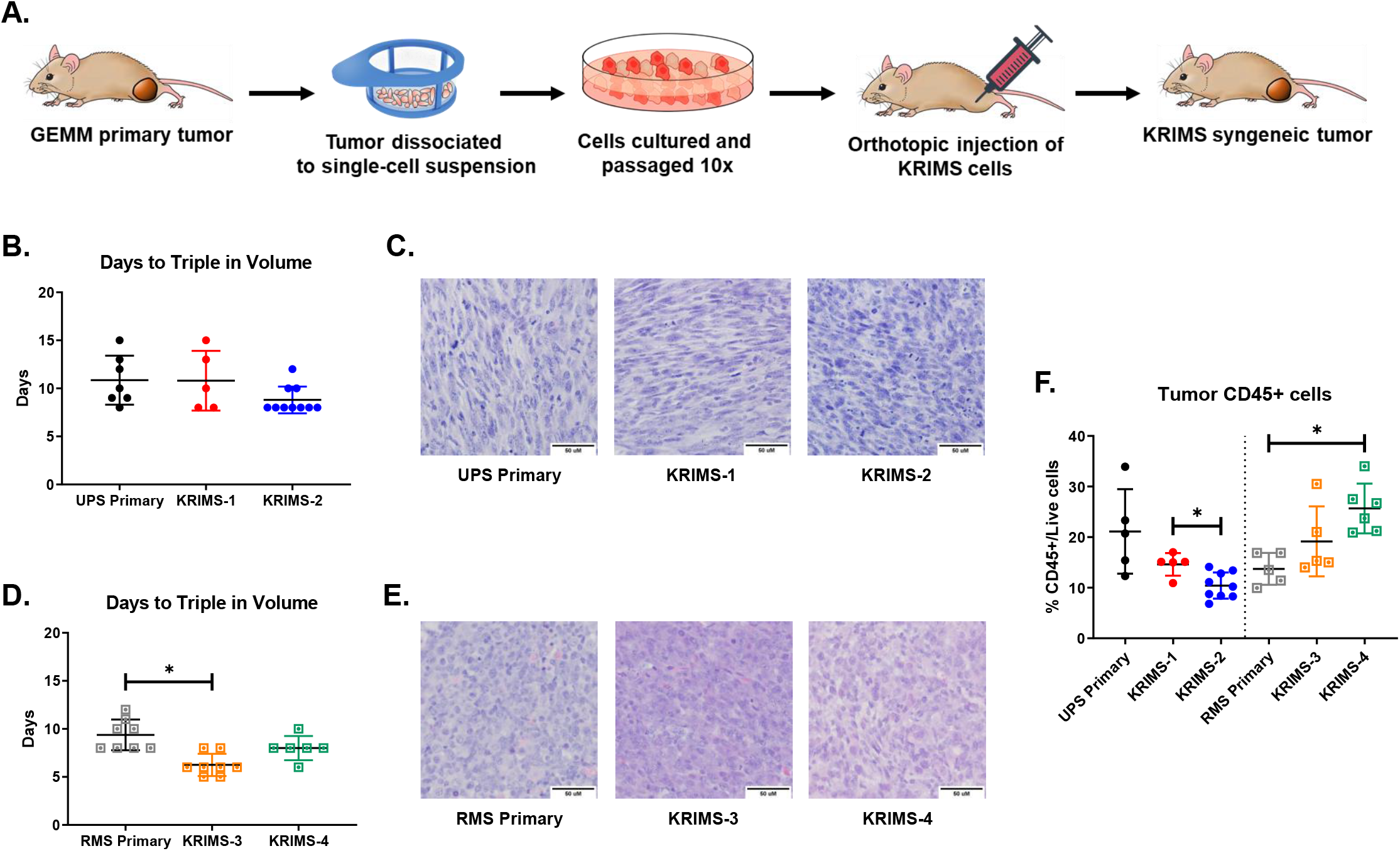
Development of orthotopic syngeneic models from UPS and RMS primary tumors. (A) Generation of syngeneic models. Expression of oncogenic *Kras^G12D^* and deletion of *Trp53* to drives UPS and RMS development in genetically engineered mouse models (GEMMs). Tumors were harvested and cells dissociated into a single cell suspension. Cells were grown *in vitro* for 10 passages to remove contaminating stroma and form <u>K</u>ras <u>I</u>nduced <u>M</u>urine <u>S</u>arcoma (KRIMS) cell lines. KRIMS lines were injected orthotopically into immunocompetent mice to form syngeneic tumors. (B & D) Both primary and syngeneic models have similar growth kinetics. (C & E) H&E staining of primary and syngeneic tumors demonstrates that syngeneic tumors have high pathologic fidelity to their respective primary tumor of origin. (F) UPS syngeneic tumors contain less CD45+ cell infiltration than UPS primary tumors, while RMS syngeneic tumors contain more CD45+ cell infiltration than RMS primary tumors.

### 3.3 Immune infiltration is decreased in UPS syngeneic models and increased in RMS syngeneic models

Though there are extensive characterizations of diverse syngeneic models across multiple tumor types and mouse background strains^5–10^, there are few studies comparing the immune profiles of orthotopic allograft models with their originating primary tumor model to determine if the syngeneic tumors accurately recapitulate the primary tumors’ intratumoral immune landscapes. We performed flow cytometry on five to nine similarly-sized orthotopic allograft tumors from each model and compared levels of CD45+ immune cell infiltration (Figure 2F). Primary and syngeneic tumors contained similar numbers of live cells except for KRIMS-4 tumors, which contained significantly more live cells than RMS primary tumors (82.0% and 66.8%, respectively) (Supplemental Figure 1A). On average, 21.1% of live cells in UPS primary tumors were CD45+. However, in the UPS syngeneic tumors, CD45+ cell infiltration was reduced by 30-50%. UPS primary tumors also had a wider range of CD45+ immune infiltration compared to UPS syngeneic tumors. Immune infiltration in RMS primary and syngeneic tumors showed the opposite trend. RMS syngeneic tumors contained 40-88% more CD45+ cells than the RMS primary tumors. Additionally, CD45+ cell infiltration in RMS primary tumors was less heterogeneous than in syngeneic counterparts. These results suggest that there are cancer type and cell line specific factors that modulate immune infiltration in syngeneic tumors, irrespective of initiating mutations or tumor location.

### 3.4 UPS primary and syngeneic tumor immune landscapes

To further appreciate the differences in tumor immune infiltration, we performed in-depth profiling of intratumoral CD45+ cells, comparing primary and syngeneic tumors within each cancer type. We performed comprehensive immunophenotyping of nine innate and adaptive cell populations: B cells, dendritic cells, natural killer (NK) cells, monocytes/PMNs, MHCII^hi/low^ TAMS, and T cell subsets (CD4+ T cells, CD8+ T cells, and Tregs). Previously published reports from multiple syngeneic tumor models have found a wide diversity of immune cell composition.^5–10^ Based on these studies and on our finding of differing levels of CD45+ immune infiltration, we hypothesized there would be broad differences in multiple immune cell populations between UPS primary and KRIMS-1 and KRIMS-2 syngeneic tumors.

Unexpectedly, we observed similar levels of several cell populations in both UPS primary and syngeneic tumors (Figure 3A). B cells, dendritic cells, and NK cells comprise similar proportions of total CD45+ cell infiltration in the UPS primary and KRIMS-1 and KRIMS-2 tumors (Figure 3B-D). In both models, myeloid cells comprise the majority intratumoral CD45+ cells. TAMs are the most abundant myeloid cell and account for 53-62% of all intratumoral immune cells. We decided to further analyze the TAM population by examining MHCII^hi^ and MHCII^low^ populations. MHCII^hi^ TAMs are mature, tissue-resident macrophages, while MHCII^low^ TAMs represent a phenotypically distinct macrophage population linked with tumor progression.^18–20^ In our observations, levels of MHCII^hi^ and MHCII^low^ TAMs are comparable between UPS primary and KRIMS-1 and KRIMS-2 tumors (Figure 3F and 3G). Additional intratumoral myeloid cells are monocytes/PMNs (CD11b+, F4/80-cells), which include myeloid derived suppressor cells (MDSCs). Monocytes/PMNs are significantly more abundant in UPS primary tumors than in KRIMS-2 tumors, and slightly more abundant than KRIMS-1 tumors. Examination of the intratumoral T cell compartment revealed further differences between UPS primary and KRIMS-1 and KRIMS-2 syngeneic tumors (Figure 4A). Levels of T cell infiltration were almost twice as high in KRIMS-1 and KRIMS-2 tumors than in UPS primary tumors (Figure 4B). Within the T cell compartment, levels of CD4+ T cells, CD8+ T cells, and Tregs varied greatly between tumor types. The T cell compartment of KRIMS-2 tumors showed the greatest differences compared to the UPS primary tumors, with less than half of level of CD4+ T cells and almost double the level of CD8+ T cells (Figure 4C-E). Additionally, the percent of CD4+ cells that are Tregs was over three times higher in KRIMS-2 tumors than in UPS primary tumors (Figure 4F). The T cell compartment of KRIMS-1 tumors also differed from UPS primary tumors, but much less dramatically. There were no differences CD4+ T cell levels, and only slight, though statistically significant, decreases in CD8+ T cells and Tregs (Figure 4C-F). Together, these data demonstrate that syngeneic tumors have a limited and variable ability to accurately recapitulate the primary tumor immune landscape, with the most notable differences present in levels of T cell subsets. To determine if the differences in the intratumoral immune landscapes were reflective of systemic immune differences, we analyzed the immune profiles of spleens from the tumor-bearing mice in the study (Supplemental Figure 2A-L). KRIMS-1 and KRIMS-2 allografts were grown in 129/SvJae mice, the background strain of KP mice, rather than in KP mice directly. Therefore, we also analyzed spleens from tumor-naïve KP and tumor-naïve 129/SvJae mice to account for any baseline differences. Several small but statistically significant differences were observed across the various cell populations in tumor-naïve KP and tumor-naïve 129/SvJae mice. These differences were also observed in spleens from KRIMS-1 and KRIMS-2 tumor-bearing mice. The only significant difference observed in both spleen and tumor tissue was an increase in KRIMS-2 splenic and intratumoral CD8+ T cells. Taken together, these data suggest that the vast majority of the differences observed in intratumoral immune landscapes are specific to the tumor microenvironment and are not reflected systemically.

**Figure 3:**
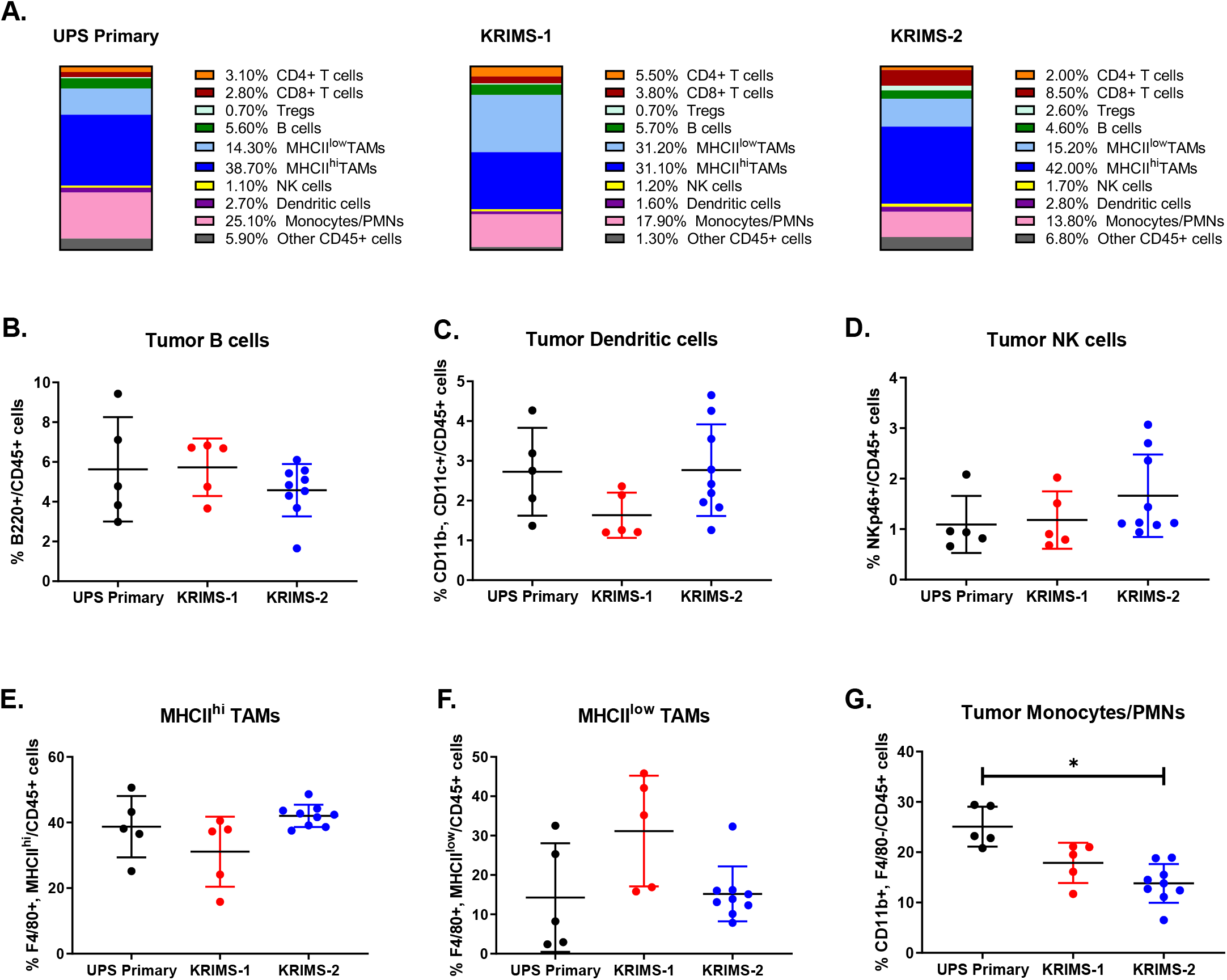
UPS primary and syngeneic tumor immune landscapes. (A) Frequencies of 9 immune cell populations, reported as percentages of CD45+ cells. Values are averages of the mice shown in the remainder of Figure 3 and Figure 4. Flow cytometry analysis demonstrating frequencies of each cell type in individual tumors including (B) B cells, (C) dendritic cells, (D) natural killer cells, (E) MHCII^hi^ tumor-associate macrophages. (F) MHCII^lo^ tumor-associate macrophages, and (G) monocytes/PMNs.

**Figure 4:**
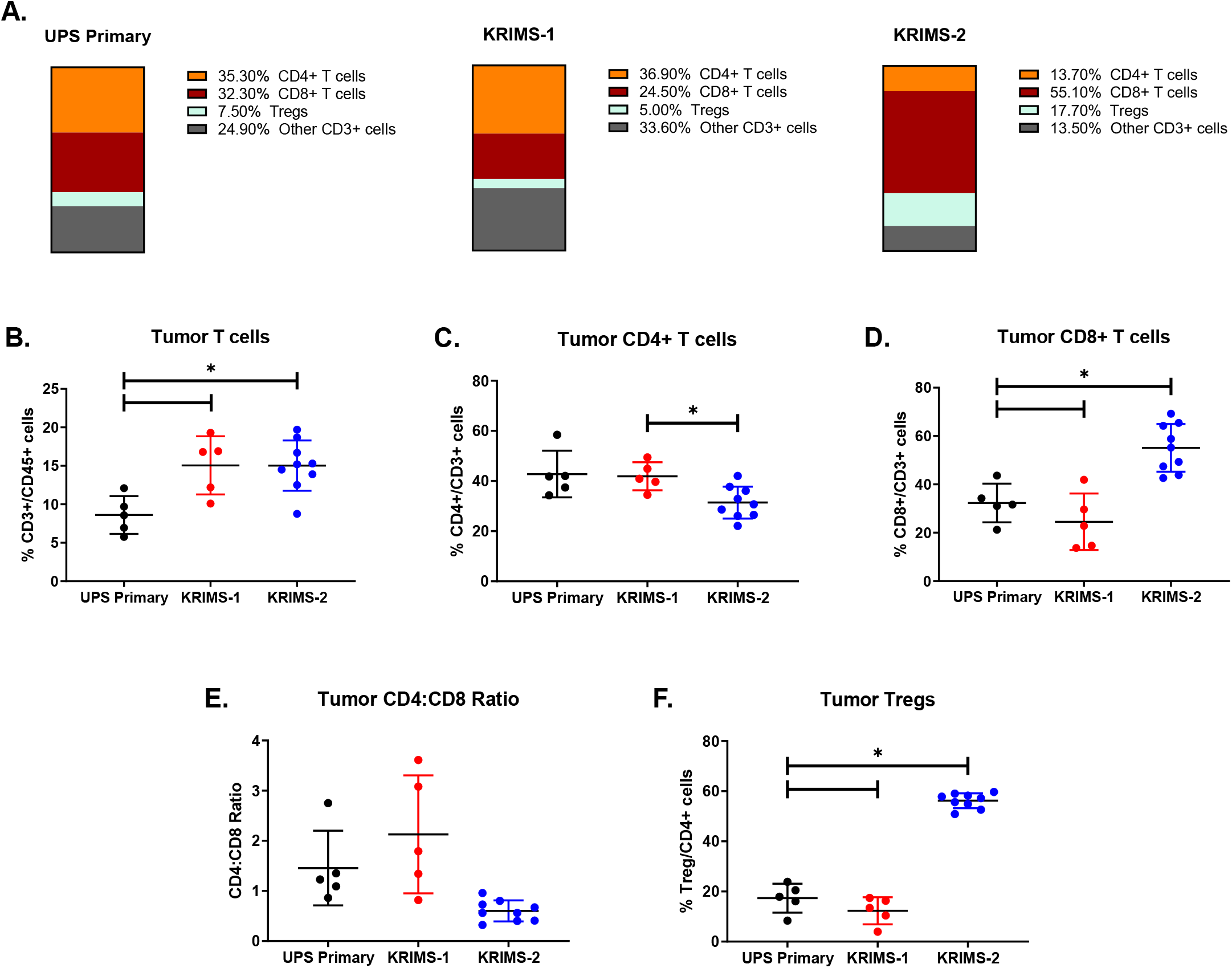
UPS primary and syngeneic tumor T cell compartments. (A) Frequencies of 3 T-cell populations, reported as percentages of total CD3+ T cells. Values are averages of mice included in Figure 3 and the remainder of Figure 4. Flow cytometry analysis demonstrating frequencies of each cell type in individual tumors including (B) CD3+ T cells, (C) CD4+ T cells, (D) CD8+ T cells, (E) CD4/CD8 ratio and (F) regulatory T cells (Tregs). Note the enriched CD3+, CD8+, and Tregs in the KRIMS-2 syngeneic model.

### 3.5 RMS primary and syngeneic tumor immune landscapes

The intratumoral immune profiles of RMS primary and syngeneic tumors showed several similarities to those of UPS primary and syngeneic tumors. Levels of B cells, dendritic cells, and MHCII^hi/low^ TAMs did not differ between RMS primary and syngeneic tumors (Figure 5A-C, E and F). Small, but statistically significant, differences in NK levels were present between KRIMS-3 and KRIMS-4 tumors, but both were within the range of RMS primary tumors (Figure 5D). Like the UPS primary tumors, one of the allograft tumors (KRIMS-4) showed a decrease in monocytes/PMNs compared to primary tumors. This difference was not statistically significant though due to the wide range of values for both sets of tumors. In contrast to the UPS tumors, levels of T cell infiltration did not differ between RMS primary and syngeneic tumors (Figure 6B). However, the composition of the T cell compartment showed significant differences and followed the same trends observed in the UPS primary and syngeneic tumors (Figure 6A). RMS primary tumors contained a significantly higher percent of CD4+ T cells, and significantly lower percent of CD8+ T cells and Tregs compared to the KRIMS-4 tumors (Figure 6C-F). KRIMS-3 tumors followed similar trends but were not significantly different from RMS primary tumors. As observed in the spleens of UPS tumor-bearing mice, several differences were observed between the three RMS tumor models (Supplemental Figure 2). However, as with the UPS tumors, the majority of these differences did not correlate with the differences in the intratumoral immune landscapes. The one exception was a slight, but statistically significant increase in Tregs in the spleens KRIMS-4 tumor-bearing mice relative to RMS primary tumor-bearing mice. However, the difference was much less robust than the two-fold increase observed in the KRIMS-4 tumors compared to RMS primary tumors. As with the UPS tumors, these data suggest that the differences in intratumoral immune cell populations are tumor microenvironment-specific. Furthermore, the data again demonstrate the limited and variable ability of syngeneic tumors to recapitulate the primary tumor immune landscape, especially within the T cell compartment.

**Figure 5:**
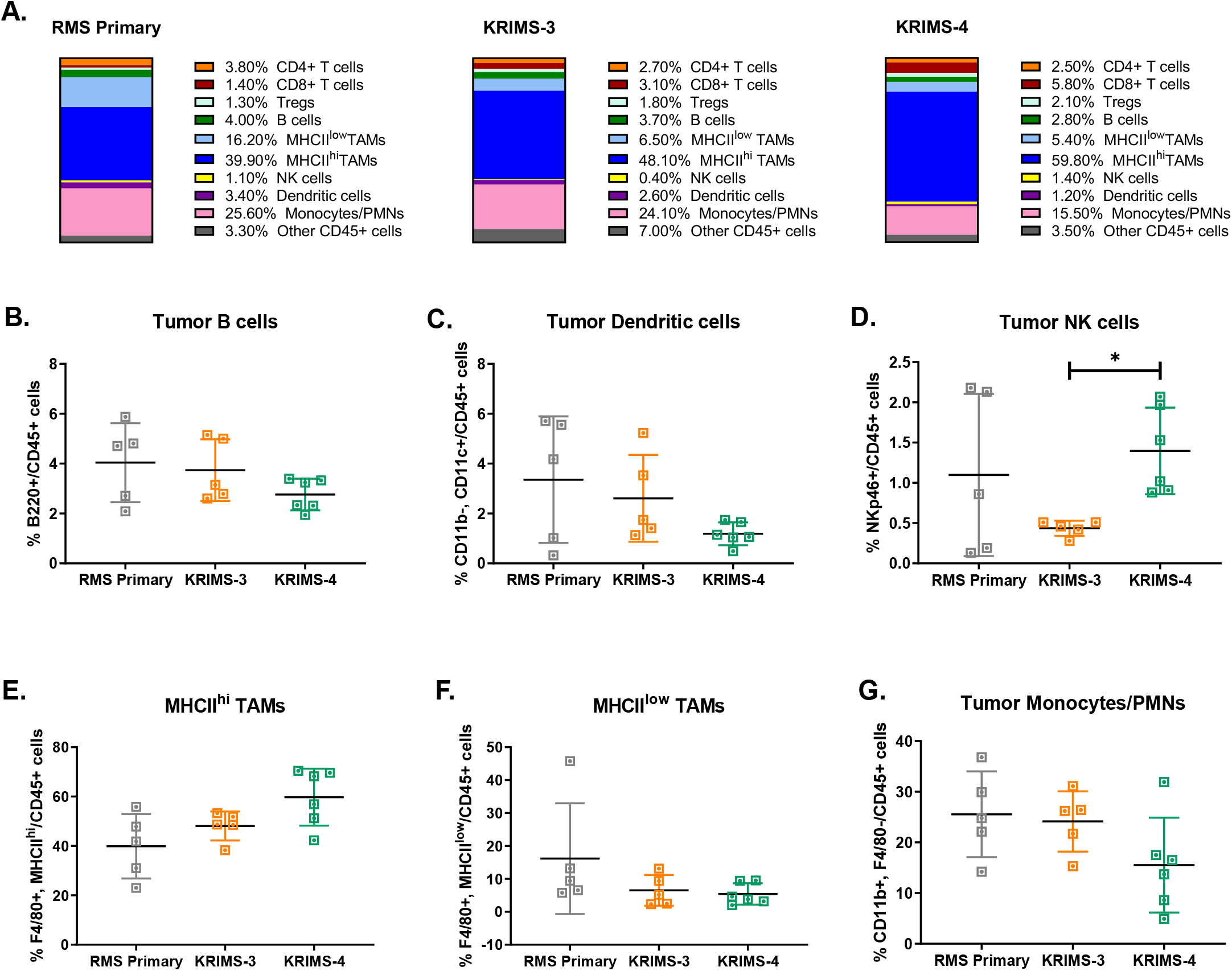
UPS primary and syngeneic tumor immune landscapes. (A) Frequencies of 9 immune cell populations, reported as percentages of CD45+ cells. Values are averages of the mice shown in the remainder of Figure 6 and Figure 6. Flow cytometry analysis demonstrating frequencies of each cell type in individual tumors including (B) B cells, (C) dendritic cells, (D) natural killer cells, (E) MHCII^hi^ tumor-associate macrophages. (F) MHCII^lo^ tumor-associate macrophages, and (G) monocytes/PMNs.

**Figure 6:**
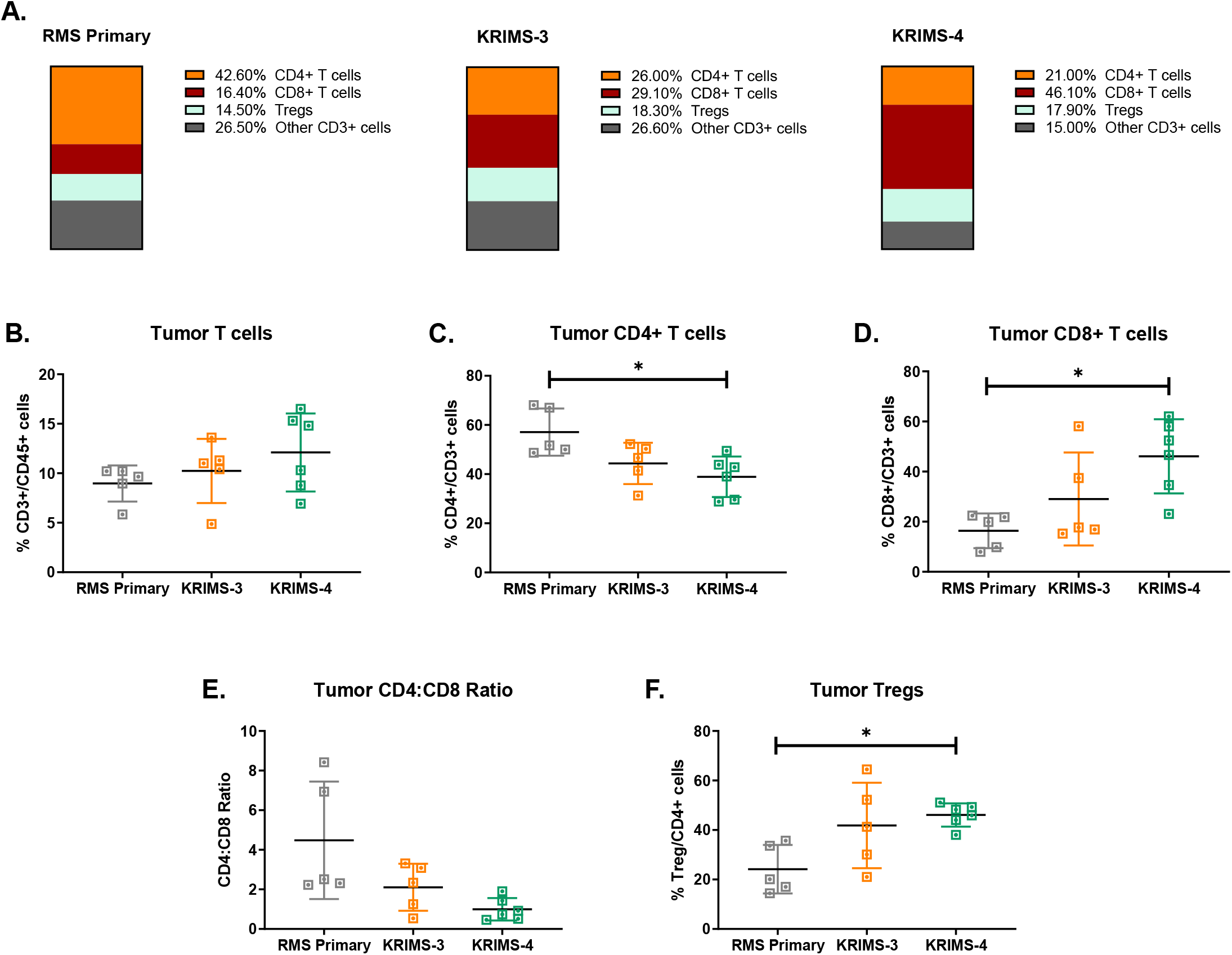
RMS primary and syngeneic tumor T cell compartments. (A) Frequencies of 3 T-cell populations, reported as percentages of total CD3+ T cells. Values are averages of mice included in Figure 5 and the remainder of Figure 6. Flow cytometry analysis demonstrating frequencies of each cell type in individual tumors including (B) CD3+ T cells, (C) CD4+ T cells, (D) CD8+ T cells, (E) CD4/CD8 ratio and (F) regulatory T cells (Tregs).

## 4. DISCUSSION

While several studies have examined the immune infiltrate of genetically engineered primary tumors, there have been few analyses comparing genetically and anatomically similar primary models of cancer. Here we report the first side-by-side characterization two primary soft tissue sarcomas that arise at the same location and that contain the same initiating mutations. Using flow cytometry, we compared levels of immune infiltration and the immune profiles of UPS and RMS primary tumors. We found that while overall immune infiltration did not differ between the two, significant differences existed in CD4+ and CD8+ T cell levels. The differences in T cell profiles point to the importance of cancer-specific signaling within the tumor microenvironment.

Though primary tumors are useful for examining therapeutic response across in a naturally developing tumor microenvironment, syngeneic models provide several advantages for preclinical studies including the speed of tumor growth, ease of experimental manipulation, and reproducibility across individual mice. Using UPS and RMS primary tumors, we generated a series of syngeneic UPS and RMS cell lines (KRIMS cells) that can be efficiently grown and studied in immunocompetent mice. Though syngeneic tumor models are commonplace in preclinical cancer research, there is a paucity of studies that assess if allografts accurately recapitulate the immune landscape the primary tumors from which they were derived. We sought to address this by performing the same flow cytometric analysis with the KRIMS syngeneic tumors as was performed with the UPS and RMS primary tumors. We found that UPS syngeneic tumors contained lower levels of immune infiltration than UPS primary tumors, while RMS syngeneic tumors contained higher levels of immune infiltration than RMS primary tumors (Figure 2). Furthermore, the T cell compartments of syngeneic tumors significantly differed from their primary tumor counterparts (Figures 4 and 6). We also observed heterogeneity between syngeneic models of the same cancer type. KRIMS-1 UPS syngeneic tumors tended to better recapitulate the immune landscape of UPS primary tumors than KRIMS-2 UPS syngeneic tumors (Figures 3 and 4). Likewise, the immune profile of KRIMS-3 RMS syngeneic tumors was more similar to RMS primary tumors than the KRIMS-4 RMS syngeneic tumors (Figures 5 and 6). Taken together, this analysis demonstrates the importance of examining immune cell diversity across distinct syngeneic series when considering allograft models for use in preclinical studies.

Our syngeneic sarcomas display several unique phenotypes in comparison to other syngeneic models. For example, the NK compartment in the UPS and RMS sarcoma models is very low (<2%), while NK cells are present at large numbers in RENCA and CT26 tumors (10-25%).^5^ While extrapolation from these comparisons must be done with caution, these observations suggest that NK-dependent approaches would be unsuccessful in sarcomas. Macrophages are the predominant myeloid cell population in our sarcomas, whereas MDSCs are the most prevalent myeloid cell type in 4T1 breast and B16F10 melanoma models.^5,6^ The proportion of T cells infiltrating a tumor can depend upon mouse background strain, as T cells represent 12-21% of total CD45+ cells in syngeneic BALB/c models (CT26, RENCA, and 4T1), but only 4-8% of total CD45+ cells in syngeneic C57BL/6 models (MC38, LL/2, and B16F10).^5^ Our sarcoma models span these ranges, with T cells comprising 8-15% of total immune cells in UPS and RMS primary and syngeneic tumors. Of note, the majority of published studies of syngeneic models report a CD4/CD8 ratio equal to or above 1.^5–10^ All our tumor models except the KRIMS-2 UPS syngeneic tumors followed this pattern. KRIMS-2 tumors, in contrast, favored CD8+ T cell enrichment with a CD4/CD8 of 0.6 (Figure 4E). Such data suggests that the best approach for studying cancer immunology may be to examine multiple syngeneic series of tumors.

In contrast to other solid tumors, immunotherapy trials in soft-tissue sarcomas have been disappointing.^21–25^ Sarcomas are a very diverse group of cancers, with over 50 distinct histological subtypes. Among sarcomas, the UPS subtype has shown the most promising responses to checkpoint blockade. Relatively few immunotherapy trials have been conducted in RMS, likely due to its predominantly pediatric incidence.^26^ In the expansion cohort of SARC028, a multicenter phase II study, UPS was the only subtype that responded to pembrolizumab (anti-PD1).^21,22^ Similar results were seen in a phase II trial for metastatic sarcoma using nivolumab (anti-PD1) with or without ipilimumab (anti-CTLA-4).^25^ In patients receiving the combination immunotherapy, only 16% (6/38) achieved a confirmed objective response. UPS was the predominant subtype among responding patients, although patients with other subtypes did respond.^25^ Current studies are attempting to increase the efficacy of immunotherapy by combining checkpoint blockade with chemotherapeutic agents (ClinicalTrials.gov identified NCT02888665) or radiation therapy (ClinicalTrials.gov identified 03092323). Since both chemotherapy and radiotherapy can increase expression of immune-related genes in cancer cells, a main objective of these approaches is to increase intratumoral immune cell recruitment.^27,28^

In an attempt to understand the limited success of immunotherapy trials and generate predictive algorithms of response, several groups have examined the immune landscape of soft-tissue sarcomas.^29–32^ In general, sarcomas are not traditionally considered “strongly immunogenic” tumors, as they contain less tumor-infiltrating lymphocytes than other solid tumors, such as renal cell carcinoma and melanoma.^33^ Our primary and syngeneic models have similarly low levels of immune infiltration, as CD45+ cells comprise only 10-20% of live cells in these tumors. In comparison to other sarcomas, the UPS subtype has the highest levels of TCR clonality, T cell infiltration, and PD-1 expression, although these levels are still low in relationship to epithelial tumors.^29,30^ B cell rich tertiary lymphoid structures, in addition to PD-1 expression, have also been shown to be key elements of response to immunotherapy in sarcomas.^34^ In our models, UPS and RMS tumors contained similar levels of B cell and CD3+ T cell infiltration, suggesting that RMS may respond to immunotherapy at similar rates as UPS.

With the increasing number of studies characterizing the sarcoma immune landscape, our knowledge of sarcoma immunology is rapidly expanding. Our studies highlight how an expanded knowledge of the immune landscape in primary and syngeneic tumors will be key to designing successful preclinical studies for sarcoma. Recently, promising combination therapy approaches that are designed to increase immune infiltration have entered the clinic. Continued translational and clinical research will be crucial to understanding the mechanisms driving intratumoral immune recruitment in order to fully realize the promise of immunotherapy for sarcoma patients.

## Author Contributions

R.D. and W.G. conceived and designed the study. W.G., A.S., G.M., E.L., and V.K.A performed the experiments. W.G. collected and analyzed the data. R.D., W.G. and A.S. interpreted the data. R.D., W.G., and A.S. wrote the paper. R.D. acquired the funding and supervised the study. All authors reviewed and approved the manuscript.

## Acknowledgments

We are grateful to personnel in the Flow Cytometry Core at the University of Iowa College of Medicine and Holden Comprehensive Cancer Center for their assistance with our experiments, particularly Justin Fishbaugh. We also thank colleagues in the Iowa Sarcoma Research Group for their critical feedback throughout this study.

