## Supplemental Figures 1-2 for "Profiling the intratumoral immune landscapes of primary and syngeneic Kras-driven sarcoma mouse models"

### Supplemental Figure 1

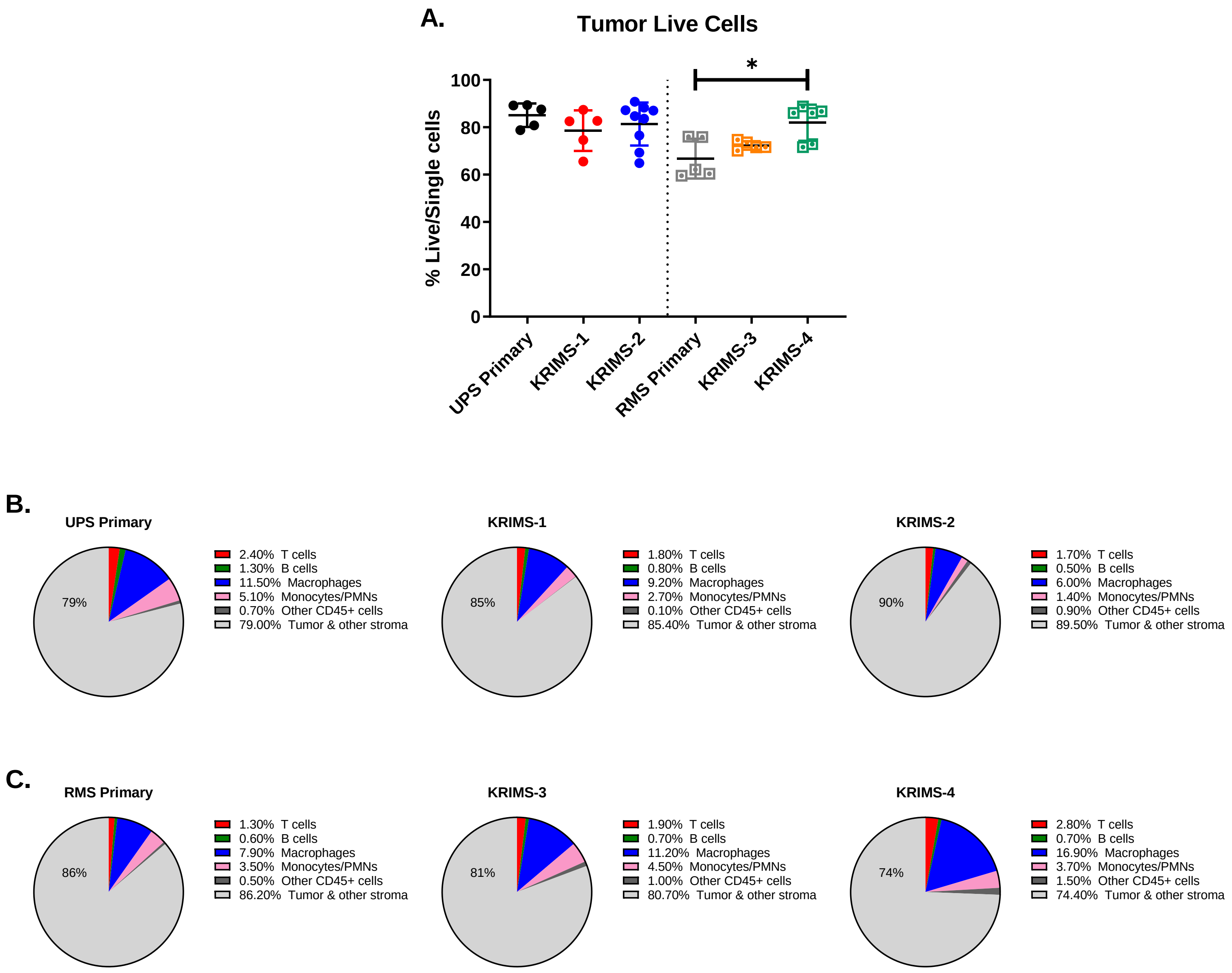


<u>Supplemental Figure 2: UPS and RMS tumor live cells and tumor composition.</u> (A) Live cells in UPS and RMS primary and syngeneic tumors. (B-C) Primary and syngeneic tumor composition as a percent of live cells. Values are an average of mice shown in (A).

### Supplemental Figure 2

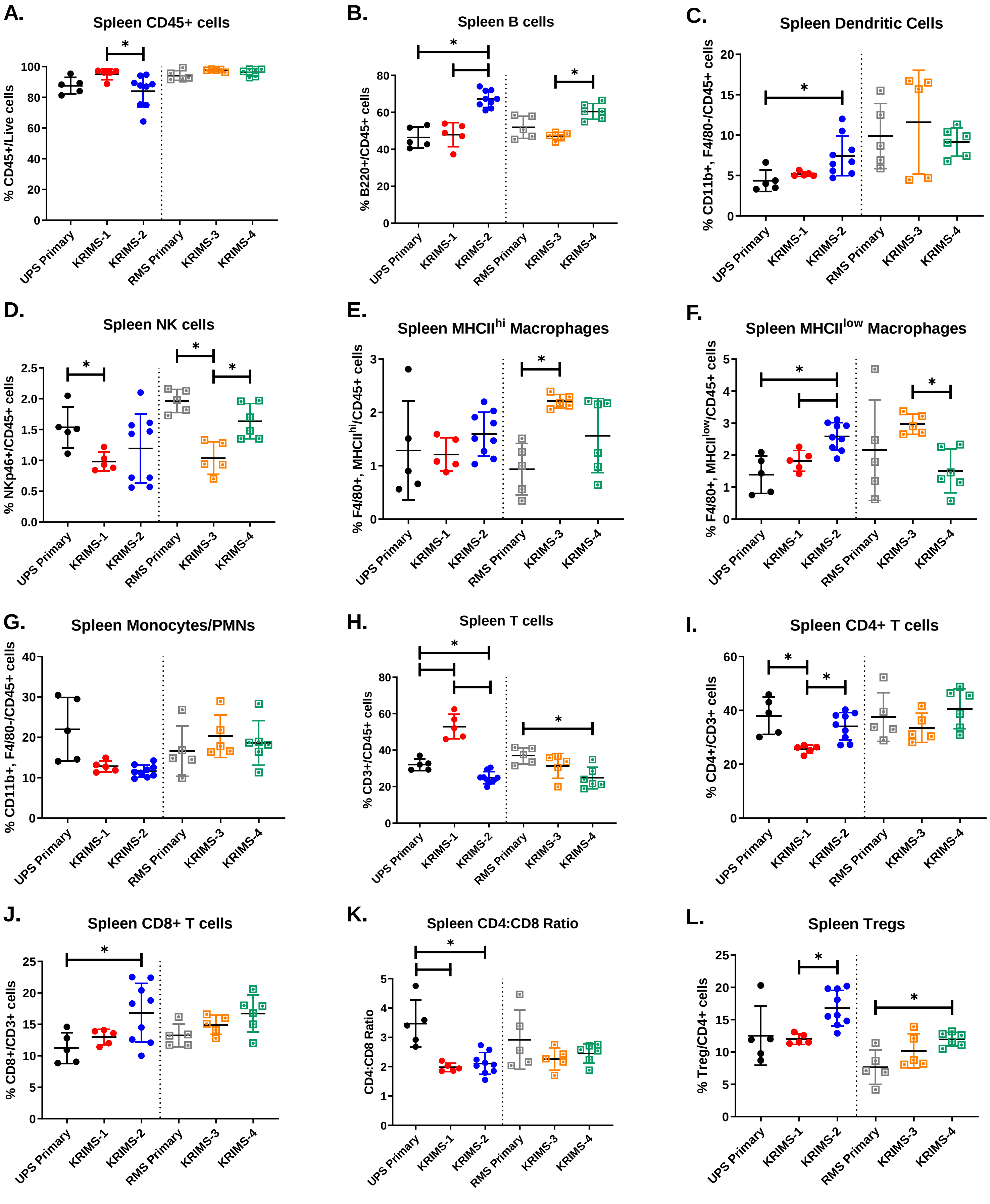


Supplemental Figure 3: Spleen immune profiles. (A-L) Immune profiles of spleens from UPS and RMS primary and syngeneic tumor-bearing mice.
